# Environmental change alters nitrogen fixation rates and microbial parameters in a subarctic biological soil crust

**DOI:** 10.1101/2022.01.25.477655

**Authors:** Alejandro Salazar, Denis Warshan, Clara Vasquez-Mejia, Ólafur S. Andrésson

## Abstract

Together, Biological Soil Crust (BSC) and other cryptogamic groundcovers can contribute up to half of the global nitrogen (N) fixation. BSC also stabilizes the soil (reducing erosion and dust emissions), fixes carbon (C), retains moisture, and acts as a hotspot of microbial diversity and activity. Much of the knowledge about how climate change is affecting the composition and functioning of BSC comes from hot arid and semiarid regions. The comparatively smaller body of research on BSC from cold and mesic environments has been primarily observational, for example along chronosequences after a glacier retreat. Few studies have experimentally investigated the effects of the environment on BSC from high latitudes. Such experiments allow unraveling of relationships at a resolution that can only be achieved by controlling for confounding factors. We measured short-term (2-4 days) responses of a liverwort-based (*Anthelia juratzkana*) BSC from the south of Iceland to a range of temperature, moisture and light conditions. Warming increased N fixation rates, especially when moisture was at a saturation level, and only when light was not limiting. A correlation analysis suggests that increases in N fixation rates were linked to cyanobacterial abundance on the BSC surface and to the rates of their metabolic activity. Warming and moisture changes also induced compositional and structural modification of the bacterial community, with consequences at the functional level. In contrast to many observations on BSC from hot drylands, the BSC from our cold and mesic study site is more limited by low temperature and light than by moisture. Our findings show possible ways in which BSC from cold and mesic ecosystems can respond to short-term manifestations of climate change, such as increasingly frequent heat waves.

## 1. Introduction

Biological Soil Crust (BSC) is a skin-like system generally dominated by cyanobacteria, fungi, lichens and bryophytes (Belnap, 2003; Bowker et al., 2018). In some areas this system can cover over 90% of the soil surface (Williams et al., 2017). BSC-forming organisms can colonize soils in harsh environments, such as drylands (Belnap, 2003; Elbert et al., 2012; Rodríguez-Caballero et al., 2018) or newly exposed soil after a glacier retreat (Breen and Lévesque, 2008; Yoshitake et al., 2018). Once they establish, they stabilize the soil (Gao et al., 2017), increase moisture retention (Breen and Lévesque, 2008), fix nitrogen (N) (Dickson 2000; Elbert et al., 2012) and carbon (C) (Yan-Gui et al., 2011; Elbert et al., 2012), and become hotspots of microbial abundance (Yoshitake et al., 2018) and diversity (Steven et al., 2013). Under constant high environmental stress, BSC becomes a permanent feature of the undisturbed ecosystem. When high abiotic stress decreases over time, BSC acts like a transient colonizer in primary succession (Bowker, 2007).

BSC covers 12% of the Earth’s terrestrial surface (Rodríguez-Caballero et al., 2018). It is estimated that before the end of the century climate change and intensification of land use will decrease this area by 25-40%, thus enhancing emissions of soil dust and reducing BSC’s contributions to the global C and N cycles (Rodríguez-Caballero et al., 2018).

The rates at which BSC influences N and C cycling depend on environmental factors such as temperature, moisture and light intensity, and on intrinsic properties of its biological components, such as the N fixing activity of cyanobacteria (Belnap, 2001) and the differential decomposing activity of bacteria and fungi (Zhao et al., 2020). Together, BSC and other cryptogamic ground covers (i.e. rock crust and bryophyte and lichen carpets) globally account for up to half of the terrestrial N fixation, and 7% of the C fixation (Elbert et al., 2012; Porada et al., 2014). Uncertainty in model projections remains high (global N fixation estimates ranging between 3.5 and 34 Tg yr^−1^; Porada et al., 2014), in part due to limited knowledge of the extent and structure of crusted communities in many ecosystems (Ferrenberg et al., 2017).

Cold-adapted BSC covers vast areas at high latitudes (Pushkareva et al., 2016), the part of the world experiencing the fastest warming (IPCC, 2021). One of the most visual and consequential manifestations of climate change in these regions and globally is the increase in frequency and intensity of short-term climatic anomalies such as heat waves (Perkins et al., 2012). Some high-latitude systems are more resistant to climatic anomalies than others (Jónsdóttir et al., 2005). Because of its relative simplicity and the fast turnover rates of its biological componentes, BSC may be particularly sensitive to short-term changes in the environment.

The potential effects of climate change on cold-adapted BSC, and on the ecosystem services it provides, are still largely unknown. Most of the knowledge on BSC responses to the environment comes from hot arid or semiarid ecosystems (e.g. Eldridge et al., 2006; Bowker et al., 2008; Büdel et al., 2009; Delgado-Baquerizo et al., 2013; Zhao et al., 2020). The comparatively smaller body of research on BSC adapted to cold and mesic conditions has been primarily observational, for example along chronosequences after a glacier retreat (Breen and Lévesque, 2008; Yoshitake et al., 2010, 2018; Borchhardt et al., 2019) or along climatic gradients (Stewart et al., 2011; Blay et al., 2017; Pushkareva et al. 2021). While observational approaches are useful to study ecosystem processes under natural conditions, experiments allow study of relationships at a resolution that can only be achieved by controlling for confounding factors. Only a handful of studies have used an experimental approach to investigate the effects of the environment on cold-adapted BSC (e.g. Colesie et al., 2014; Alatalo et al., 2015; Rousk et al., 2018).

From a methodological perspective, experimental research on BSC offers many advantages (Bowker et al., 2014; Maestre et al., 2016). Because of its size, it is possible to collect entire BSC blocks and study them as closed systems, even under laboratory conditions (Figure S1). Also, because of the small size and relatively fast turnover rates of BSC organisms, it can respond rapidly to external factors. This has advantages for early detection of environmental changes. Finally, since BSC can act as the first link in a chain of ecological succession (Godínez-Alvarez et al., 2011), understanding BSC responses to the environment not only provides information about the BSC itself, but also about the potential ecosystems and ecological dynamics that can evolve from it.

Here we conducted a laboratory experiment to study the compositional and functional responses of a cold- and mesic-adapted BSC to short-term (2-4 days) incubations at a range of temperature, moisture and light intensities. The ranges of temperature varied between average and maximum values during the snow-free season in an Icelandic ecosystem dominated by BSC. Unless limited by other biotic and/or abiotic factors, we expected warming to increase N fixation rates either via growth of N fixers, or via increases in their metabolic activity, or both. Also, we expected environmental treatments to exert a selective pressure, leading to structural and possibly compositional changes within the N fixing communities (e.g. favoring warm adapted taxa); and for this to contribute to explaining N fixation responses to the environment. Given the well known dependence of biological N fixation on moisture (Rousk et al., 2018), and the need of photosynthetically synthesized C to run the N fixing machinery (Scherer et al., 1988), we expected the effects of warming on N fixation to be directly dependent on moisture and light intensity. Also, we hypothesized a positive correlation between N fixing rates and chlorophyll *a* (Chl *a*) content - an indicator of net photosynthetic rate (Yan-Gui et al., 2011). Overall, we expected subarctic BSC to respond to short-term environmental manipulations and for these responses to indicate possible ways in which this system, and the key ecosystem services it provides, could be altered by climate anomalies like increasingly frequent heat waves (Perkins et al., 2012).

## 2. Materials and methods

We designed a controlled laboratory experiment to investigate the responses of subarctic BSC from the south of Iceland to different levels of temperature, moisture and light. We studied how these environmental factors affect the capacity of subarctic BSC to fix N, and whether these responses were linked to changes in the abundance of N fixers and/or to structural changes in the BSC microbial communities.

### 2.1. Sample collection

In September 2018 we collected BSC from a site adjacent to the *Climate Research Unit at Subarctic Temperatures* (CRUST) experiment (Salazar et al., in progress), near Landmannahellir, Iceland (64°02’ N, 19°13’ W; 590 m.a.s.l.). Mean annual temperature and precipitation at the site are ca. 5 °C and 1500 mm, respectively. Surface cover in this area is primarily liverwort-based BSC (ca. 50%), followed by mosses (ca. 30%) and *Salix herbacea* dwarf willow (ca. 20%), on an andosol/vitrisol substratum.

We randomly collected eight BSC blocks (i.e. replicates) of 13×16 cm^2^ and ca. 5 cm deep (Figure S1a). Blocks were separated by at least 10 meters. Since the focus of this study is on BSC, patches of moss or vascular plants were avoided. We transported (approx. 5 h) the BSC blocks in coolers with ice packs and stored them in a dark room at 5 °C for 2 to 5 weeks while we performed the analyses described below. We kept wet paper towels inside the coolers to prevent desiccation.

### 2.2. Experimental design and environmental treatments

We studied the effects of temperature, moisture and light on N fixation and the microbial community structure. For this, we conducted a factorial experiment (4 × 2 × 2) with four levels of temperature: 10, 15, 20 and 25 °C; two levels of moisture: ca. 75% (close to moisture at the moment of sampling) and 100% (saturated); and two levels of light ca. 2 μmol m^-2^ s^-1^ (low intensity) and ca. 90 μmol m^-2^ s^-1^ (high intensity; Figure S2). Light was available all the time (i.e. we did not set day/night cycles), to simulate conditions in the sampling site during the summer. Temperature and light treatments were set in a growth chamber (Termaks series 8000, Bergen, Norway), and monitored hourly with temperature/light loggers (HOBO Pendant® MX Temperature/Light Data Logger, MX2202, Onset, Bourne, MA, USA). Levels of these environmental variables were selected within ranges commonly experienced by BSC at the sampling site (unpublished observations) and comparable ecosystems (Rousk et al., 2018).

Average temperature and light intensity inside the jars were 11.1 ± 0.7, 16.5 ± 0.7, 21.5 ± 0.7 and 26.6 ± 0.9 °C (2 loggers x 2 light levels; n = 4) and 2.3 ± 0.04 and 88.0 ± 1.6 μmol m^-2^ s^-1^ (2 loggers x 4 temperature levels; n = 8) respectively (Figure S2a and b). Temperature levels inside the jars were slightly higher than temperatures set in the growing chamber due to a greenhouse effect.

To create a saturation level in the moisture treatment, we wetted each sample with an excess of deionized water and waited for approximately one minute until it stopped dripping. Moisture was maintained between analyses by placing wet towels in the coolers stored in the cold, dark room. After environmental treatments and N fixation measurements (see following section), we oven dried (60 °C, 24 h) BSC disks to estimate the dried weight of the samples, and to prepare them for chlorophyll *a* analysis and DNA extraction. Average moisture content was 75.5 ± 2.4 and 107.2 ± 2.3 % (Figure S3c).

### 2.3. N fixation under controlled temperature, moisture and light conditions

We estimated N fixation rates with the Acetylene Reduction Assay (ARA; Hardy et al., 1968). We subsampled disks of 5 cm diameter and 1.5 cm depth (Figure S1c) out of the 13×16×5 cm^3^ BSC blocks (Figure S1a). We used eight 5 cm-diameter disks (i.e. replicates) per combination of temperature and moisture treatments. Thus, each temperature-specific ARA analysis was composed of a total of 16 samples with two levels of moisture, eight saturated and eight unsaturated, plus controls with acetylene, ethylene and air. The BSC disks were weighed (for further water content analysis) and placed in 350 mL glass jars with rubber septa in the lids (Figure S1c). These jars were then placed in an environmental chamber (Termaks series 8000, Bergen, Norway) at fixed temperature and light conditions. We acclimated the samples to each combination of temperature and light for 24 h. We then manually aerated the jars for a few seconds, closed the jars tightly and replaced 10% of the headspace with acetylene (except in jars used as ethylene and air controls). We incubated the jars at the set temperature and light conditions for 24 h. Then, we collected 22 mL of gas from each jar and analyzed it using a Clarus 400 gas chromatograph (PerkinElmer Ltd., Beaconsfield, UK) equipped with an automatic split/splitless injector and a flame ionization detector (FID), and an Elite-Alumina column (30 m, 0.53 mm; PerkinElmer Ltd., Beaconsfield, UK).

At the end of each 48 h acclimation-incubation period, we manually aerated the samples and started a new acclimation-incubation at a different light (but same temperature) condition. To control for a possible cumulative effect between light levels, we switched the order of the light levels for each temperature treatment i.e. half of the times starting with low light, and the other half starting with bright light.

### 2.4. Cyanobacteria and liverwort cover on BSC

We estimated the cover of cyanobacteria and liverwort (*Anthelia juratzkana*; Figure S1b) on the BSC surface by epifluorescence microscopy (Figure S3). After ARA measurements, BSC samples were stored in a dark room at 5 °C for 1 to 4 days. Plant and cyanobacterial growth was assumed to be minimal under these conditions. From each 5 cm diameter BSC disk (Figure S1c), we subsampled a 1.5 cm diameter BSC disk and imaged the plant (liverwort) chlorophyll using a Leica DM6000B fluorescent microscope (Leica, Heerbrugg Switzerland) equipped with an I3 filter cube (Ex 450/90, Di 510, Em 515), and the cyanobacterial phycocyanin with a TX2 filter cube (Ex 560/40, Di 595, Em 630/30). Multiple fields of view were measured using both filter cubes and stitched together to form an image of 1×1 cm of BCS surface (Figure S3) using the Leica software. Images were analyzed in ImageJ/Fiji (Collins, 2007; Schindelin et al., 2012), and estimates of cyanobacterial and plant covers calculated as percentage of BSC surface cover.

We did not subsample BSC disks between light levels, but rather used samples that were exposed to low light for 48 h (24 h acclimation plus 24 h ARA) and then to high light for another 48 h, or vice versa. Therefore, the treatments in this part of our analysis include temperature and moisture, but not light.

### 2.5. Chlorophyll *a*

We estimated Chl *a* content as an indicator of net photosynthetic rate in BSC (Yan-Gui et al., 2011). Similar to our BSC cover analysis, we subsampled a 1.5 cm diameter BSC disk from each 5 cm diameter BSC disk (Figure S1c) used for ARA analysis. We dried subsamples at 60 °C for 24 h, extracted Chl *a* using DMSO (65 °C, 90 min) and then estimated Chl *a* content by spectrophotometry (665 and 750 nm; Genesys 20, Thermo Scientific, Waltham, MA), as in Caesar et al. (2018):

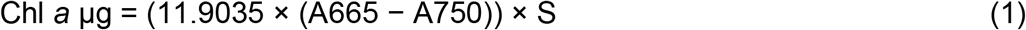

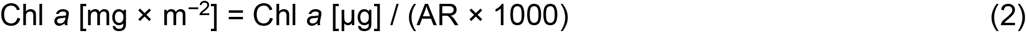

Where S is volume of solvent, AR is area (in m^-2^) and A665 and A750 are absorbances at 665 and 750 nm, respectively.

As for BSC cover, treatments in this part of our analysis included temperature and moisture, but not light.

### 2.6. DNA extraction and analysis

Immediately after the fluorescence microscopy measurements (section 2.4), we dried (60 °C, 24 h) and ground (1 min, Mini bead beater 16; Biospec products) the 1.5 BSC disks used for the cyanobacteria/liverwort cover analysis and stored them at -80 °C for up to four months for DNA extraction. We pooled together replicates in pairs, combining them in equal weight parts (125 mg each for a total of 250 mg). We used the PowerSoil® DNA extraction kit (MOBIO/Qiagen), and shotgun sequencing approaches and analyses via the alignment-free fast taxonomic annotation tool Kraken2 (Wood and Langmead, 2019) with the Kraken2 Refseq Standard plus protozoa and fungi database and the web-based pipeline Kaiju (Menzel et al., 2016). We estimated relative abundance of microbial groups using Kraken2 and fungal:bacteria ratios based on Kaiju taxonomic assignments (see sections below). After quality filtering the raw reads using Trim Galore microbial metagenome functional profiling was performed using HUMAnN 3 (Beghiji et al., 2021). For the functional annotation, UniRef50 (Suzek et al., 2015), KEGG (Kanehisa and Goto, 2000), and BioCyc databases (Karp et al., 2019) were used. As for BSC cover and Chl *a*, treatments for this part of our analysis included temperature and moisture but not light. We characterized microbial communities only at two temperature levels: 10 and 20 °C, which showed significant differences in N fixation and cyanobacterial cover (see Results).

### 2.7. Fungal:bacterial ratios

Fungi and bacteria decompose organic matter at different rates, which affects the N and C biogeochemistry of substrates like BSC. To study potential effects of the environment on the biogeochemistry of BSC via differential effects on fungi and bacteria, we estimated fungal:bacterial ratios. We calculated fungal:bacterial ratios based on numbers of gene copies assigned to each group by Kaiju.

### 2.8. Microbial community and statistical analyses

Microbial community analyses were performed using the microeco package in R (version 3.5.0). We first investigated the most important Orders for classifying samples into different treatments using a random forest approach. We then conducted an ANOVA test followed by a Tukey’s HSD test, α<0.05, as well as Pearson correlations and PERMANOVA analyses between the Bray–Curtis dissimilarity score and moisture content. Finally, we conducted a Distance-based redundancy analysis (dbRDA) to assess the effects of the abiotic treatments on the top most abundant bacterial orders. To identify distinctive molecular pathways between treatments, we performed a linear discriminant analysis (LDA) effect size (LEfSe) analysis as implemented in the microeco package, then we selected the functions with a LDA score ≥ 3.5.

We used a mixed model (*lmer* function in R, version 3.6.1) to analyze the fixed effects of environmental manipulations on N fixation, while accounting for the random effect of measurements on the same sample at two light levels. For the other response variables, which varied in response to temperature and moisture but not light, we used fixed models (*lm* function in R, version 3.6.1). We compared models based on the Bayesian Information Criterion (BIC; Figure S4).

## 3. Results

### 3.1. Nitrogen fixation potential

N fixation (using acetylene to ethylene conversion as a proxy) in BSC increased with light, temperature and moisture (Figure 1). Responses to environmental manipulations were better explained by the direct and additive effects of light, temperature and moisture than by their interactions (FigS4a). The minimum N fixation (ca. 0.5 nmol cm^-2^ h^-1^) occurred in the BSC at low temperatures, regardless of moisture and light intensity, whereas the maximum (4.4 ± 0.5 nmol cm^-2^ h^-1^) occurred in moisture saturated BSC at 25 °C and high light intensity (Figure 1).

**Figure 1.**
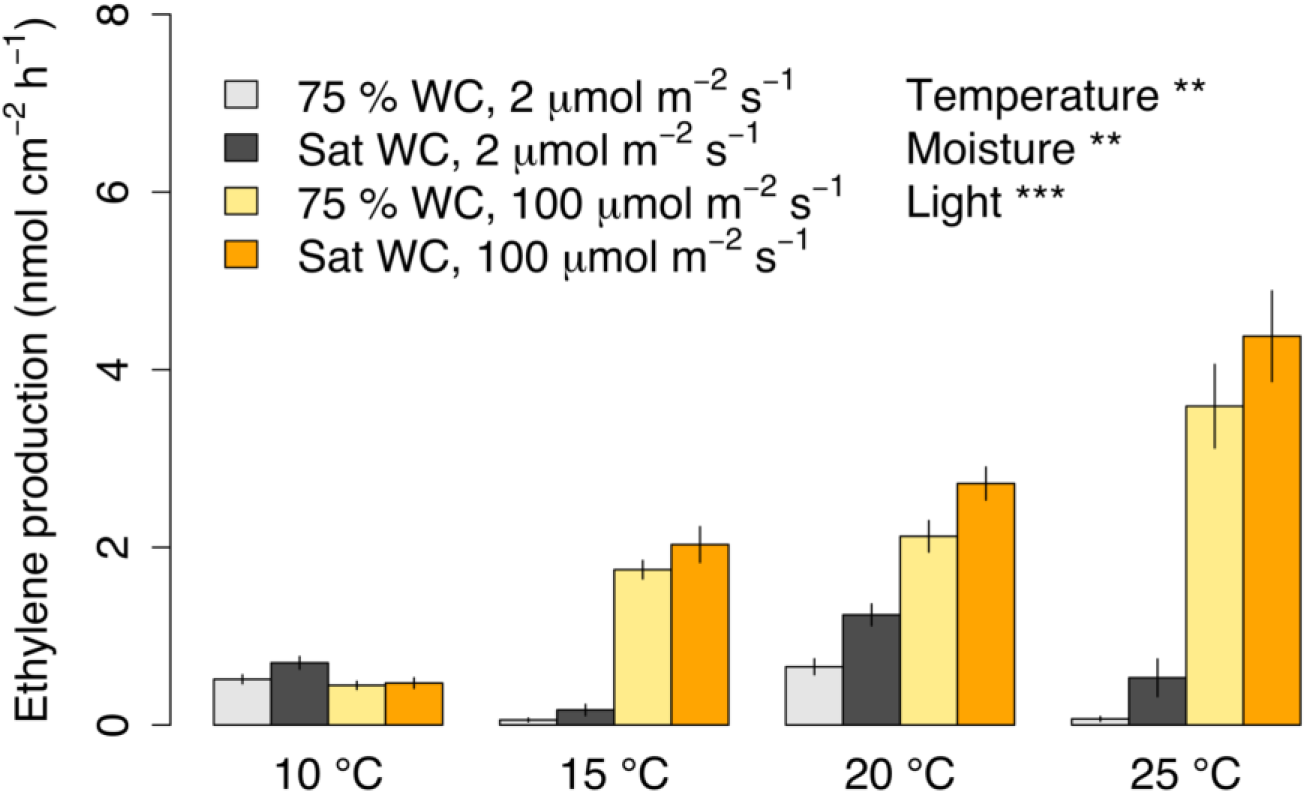
Ethylene production in response to temperature, moisture (light and dark tones), and light intensity (grey and yellow colors). Significance codes (here and elsewhere): P < 0.001 ‘***’, P < 0.01 ‘**’. Significance levels shown in the legend are from the statistical model with the lowest BIC in Figure S4a. Values are means ± se. n = 8. WC: Water content. Sat: Saturated.

### 3.2. Cyanobacteria and liverwort cover

Cyanobacterial cover on BSC increased with temperature (p<0.05) and was not affected by moisture (Figure 2). BSC incubated at 20 °C had ca. 5 to 30% more cyanobacteria on the surface than BSC incubated at 10 °C. There was no difference in cyanobacterial cover between BSCs incubated at 20 and 25 °C Liverwort cover was not affected by temperature or moisture (Figure 2). Temperature alone was a better predictor of cyanobacterial cover than moisture, or temperature and moisture combined (Figure S4).

**Figure 2.**
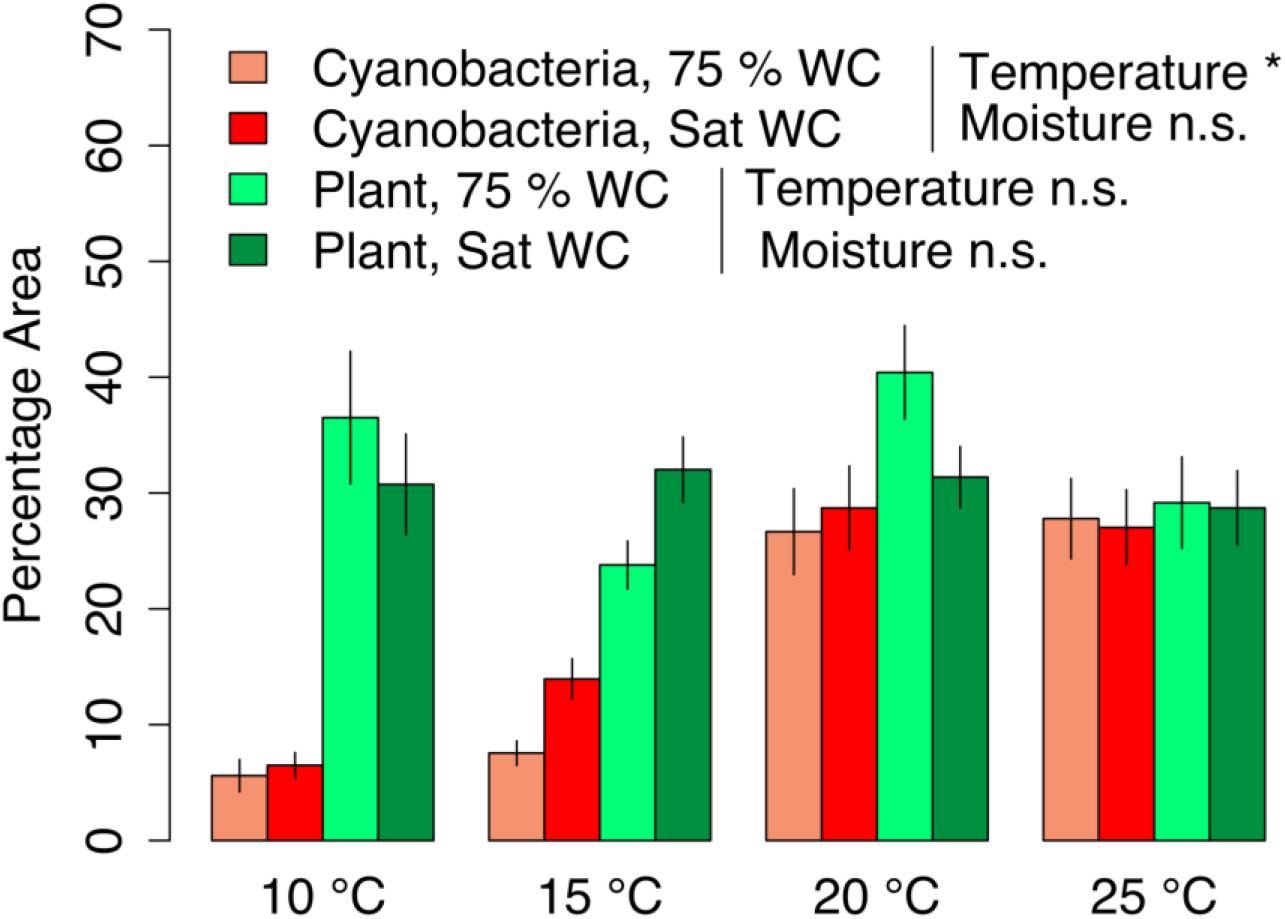
Cover of cyanobacteria and liverwort after 96 hours incubation under different conditions. WC: water content (%). Significance codes (here and elsewhere): P < 0.05 ‘*’. Values are means ± se. n = 8. WC: Water content. Sat: Saturated.

### 3.3. N fixation and cyanobacterial cover

Increases in N fixation rates were correlated (p < 0.05) to cyanobacterial cover (Figure 3). Both cyanobacterial cover and N fixation rates increased with temperature between 10 and 20 °C. Between 20 and 25 °C N fixation rates continued increasing (Figure 1) but cyanobacterial cover did not change (Figure 2). Chl *a* did not vary with environmental treatments (Figure S5) and was not related to N fixation rates (Figure S6).

**Figure 3.**
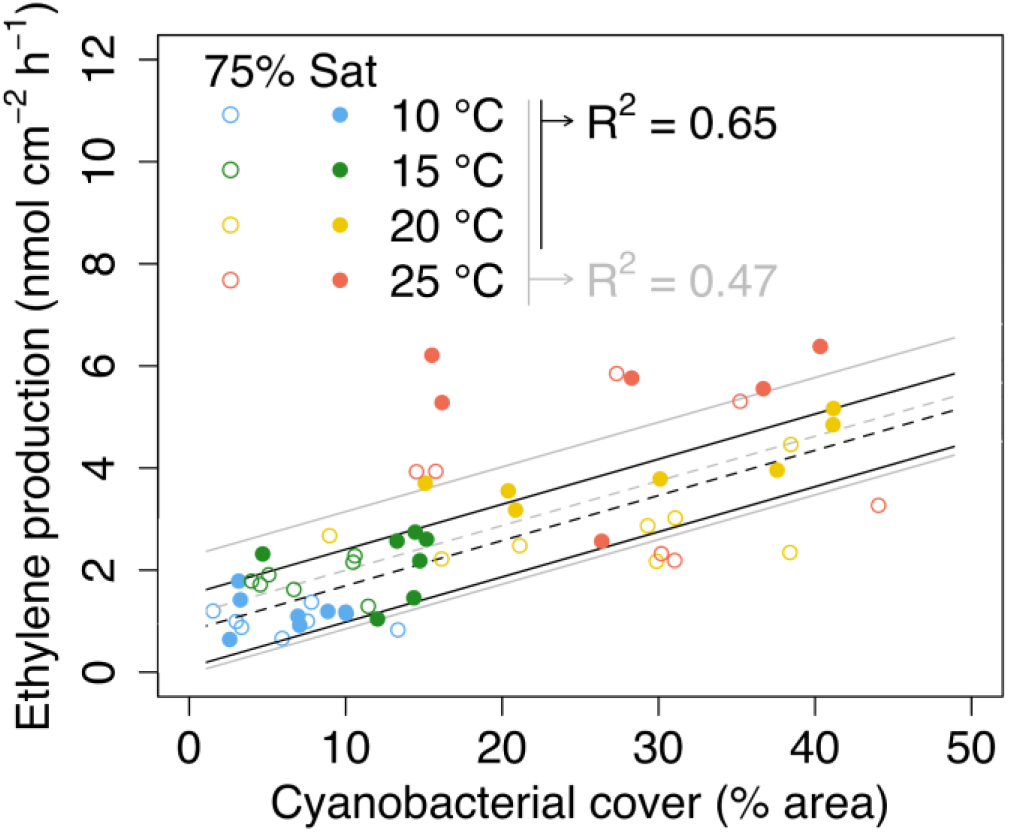
Correlation between cyanobacterial cover and N fixation rates across different temperature, moisture and light levels in a subarctic BSC. Dashed lines show the mean regression line, and solid lines the standard deviation of the mean. Grey lines and text show correlation between cyanobacterial cover and ethylene production for all temperature treatments (*y*_*i*_ *= 1*.*12 + 0*.*09*x*_*i*_ *+* □; □∼*N(0, 1*.*16*^*2*^), where □, *here and elsewhere, is the error term describing the random component of the linear relationship*). Black lines and text, show results excluding the 25 °C level (*y*_*i*_ *= 0*.*81 + 0*.*09*x*_*i*_ *+* □; □∼*N(0, 0*.*74*^*2*^)—to illustrate the strong correlation between cyanobacterial cover and ethylene production between 10 and 20 °C. P < 0.05 ‘*’, with and without including the 25 °C level. (n=8). Sat: Saturated (water content).

### 3.4. Relative abundance of cyanobacteria

Of the 20 most abundant Orders in the BSC microbial community, six had the highest importance score given by the random forest classification algorithm (Figures 4a and b and S7). Alphaproteobacteria belonging to the Burkholderiales and Sphingomonadales, as well as Actinobacteria from Corynebacteriales were more abundant at 20 °C than 10 °C; Burkholderiales and Sphingomonadales being more abundant at 20 °C with saturated water content (Figures 4a and b). Cyanobacteria from the Nostocales are overall more abundant at 10 °C than 20 °C. Actinobacteria from Pseudonocardiales and Planctomycetes from Isosphaerales are more abundant at 75% water content than in saturated BSC. Pearson correlations and PERMANOVA analyses between the Bray–Curtis dissimilarity score and moisture content showed a significant effect of moisture on the microbial community structure (Figure 4b, Table S1). Moisture had a significant effect on the α-diversity, based on the observed richness and Chao1 index of microbial taxa at the phylotype level (Figure 4c, Table S2 and S3). Significant differences in Chao1 were observed between 75% and saturated BSC at 10 °C and at 20 °C, with overall higher richness observed in saturated BSC than 75% (Figure 4c).

**Figure 4.**
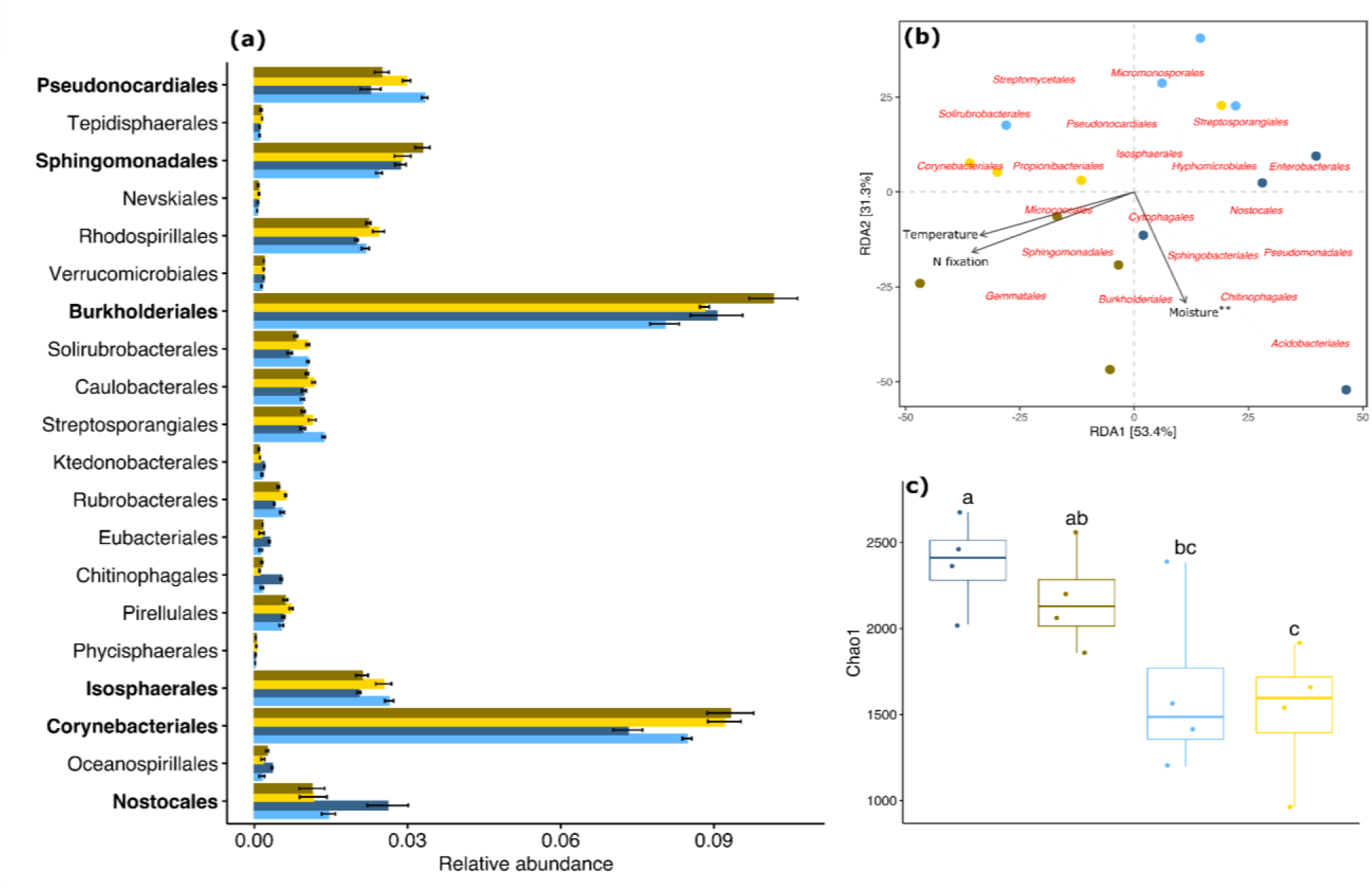
Microbial community composition and richness in a subarctic BSC under different temperature and moisture treatments. Blue and yellow indicate 10 and 20 °C, and light and dark tones indicate 70 and 100% moisture content, respectively. a) Relative read abundances of the 20 most important Orders in our study system, identified by Random Forest classification (see Figure S7). Taxa belonging to the 20 most abundant Orders in the BSC are highlighted in bold. b) dbRDA ordination plots of the community-treatment relationships of microbial communities. The top 20 most abundant Orders are shown in red. The stars (r=0.30, P < 0.01) represent significant Pearson correlations between the Bray–Curtis dissimilarity score and moisture content using mantel test. c) The α-diversity Chao1 richness index across treatments. Different letters above data points indicate statistically significant differences (P<0.05).

Fungal:bacterial ratios were slightly affected by a combination of warming and wetting (Figure S8). Total gene numbers indicate that bacteria grew faster than fungi in the saturated BSC at 20 °C, resulting in a decrease in fungal:bacterial ratios (Figure S7). Total DNA did not change significantly across environmental treatments (P>0.05), but it tendended to increase with moisture, especially under warming (Figure S8).

### 3.5 BSC microbial functions

The LEfSe results show the effect size of the top 15 differentially abundant MetaCyc metabolic pathways between temperature (Figure 5a) and moisture (Figure 5b). At 10 °C, pathways linked to autotrophy and anaerobic respiration such as nucleoside and nucleotide biosynthesis/degradation, cofactors, carrier and vitamin biosynthesis, glycolysis and photosynthesis were significantly enriched. At 20 °C, central metabolism pathways associated with cellular respiration and fermentation were more abundant such as the pentose phosphate pathway, TCA cycle and electron transport chain. Pathways associated with fermentation and TCA cycle were found enriched at 75% water content compared to saturated water content which was characterized by an enrichment in cellular respiration related pathways as well as carbohydrate, vitamin, nucleotide, fatty acid and lipid biosynthesis.

**Figure 5.**
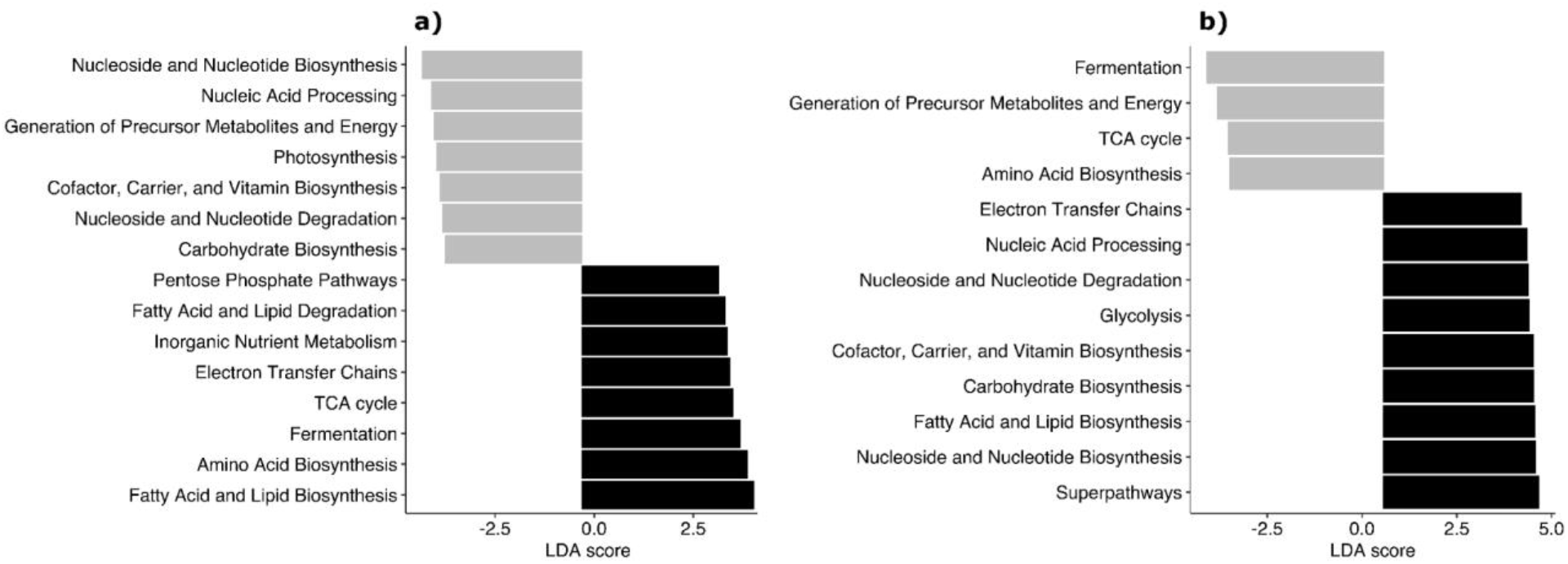
Top 15 MetaCyc pathways to be differentially abundant at a) temperature (10 °C in grey vs 20 °C in black) and b) moisture (70% in grey vs 100% in black). LDA scores were calculated using LEfSe analysis.

## 4. Discussion

Climate-driven shifts in the cover, abundance, composition and activity of BSC have the potential to alter the biogeochemistry of terrestrial ecosystems (Elbert et al., 2012; Porada et al., 2014; Weber et al., 2015; Ferrenberg et al., 2017). In hot drylands, warming alone may have little or no effect on the bacterial communities of BSC, wetting alone can increase cyanobacterial abundance, and wetting and warming combined can decrease it (Steven et al., 2015). This shows how bacterial communities from dryland BSC can buffer moderate levels of warming to some extent, but that when combined with altered precipitation the effects of warming can be noticeable (Steven et al., 2015). In our experiment with a BSC adapted to cold and mesic conditions, wetting did not have an effect on the surface cover (a proxy for abundance) of cyanobacteria, and warming increased it. This suggests that the growth of cyanobacteria in our system may be more limited by low temperatures than by lack of moisture. Alternatively, this may reflect the wider range of temperature than moisture levels in our experimental design. Mean annual temperature and precipitation are projected to increase in the high north in the coming decades (IPCC, 2021). If the Icelandic BSC used in this study is representative of BSC from cold and mesic climates in other high latitudes, our results suggest that the abundance of cyanobacteria in this type of BSC could be affected more by warming than by increased precipitation.

The time-scales at which BSC responds to the environment varies with stages of ecological succession (Pushkareva et al., 2017). In the Arctic, temperature and light intensity can more rapidly affect N fixation rates in less-developed BSC dominated by cyanobacteria than in more-developed BSC with dense lichen covers (Pushkareva et al., 2017). The BSC used in this study has the signatures of an early successional stage, including abundant cyanobacteria, being virtually free of lichen patches, having fungal:bacterial ratios closer to those found in deserts than in the tundra (Fierer et al., 2009), and having a mean Chl *a* content closer to those reported for cyanobacterial soil crust than for lichen crusts (Wu et al., 2017). Ecosystems dominated by BSC similar to the one used in this study could be particularly sensitive to climate change.

Our correlation analysis suggests that the positive relationship between cyanobacterial cover and N fixation in our study system is in part related to net abundance of cyanobacteria on the soil surface, and in part to the level of their metabolic activity. Between 10 and 20 °C, warming increased both cyanobacterial cover and N fixation, suggesting that the increase in N fixation within this temperature range may have been caused by an increase in the number of heterocysts — the N fixing cells of cyanobacteria. If total DNA is assumed to be an indicator of microbial biomass (Semenov et al., 2018; but see Leckie et al., 2004), the previous interpretation is challenged by the lack of difference in total DNA between environmental treatments. However, it is plausible that warming caused a growth of cyanobacteria and a reduction of other microbial groups, resulting in no change in total DNA. A comparison of long-term climatic experiments in drylands in the USA, for example, showed that wetting BSC increased the abundance of cyanobacteria while reducing the abundance of other photosynthetic organisms (Steven et al., 2015).

Alternatively, warming may have altered the microbial composition of the BSC profile, which varies with depth (Maire et al., 2014). Many cyanobacteria commonly found in BSC such as *Microcoleus vaginatus* are motile (Campbell, 1980), and can use this motility to reach out for light (Biddanda et al., 2015; Schuergers et al., 2016). In a laboratory experiment with cultured *Oscillatoria*-like cyanobacteria, warming increased the speed at which cells moved towards light (∼50 μm min^-1^ at 10°C and ∼215 μm min^-1^ at 35°C; Biddanda et al., 2015). An increase in cyanobacterial cover with no change in total DNA could have been caused by a temperature-accelerated migration of cyanobacteria to the BSC surface, where there is more light. There, they could have fixed more N than in deeper and darker BSC layers. Overall, our results suggest that between 10 and 20°C, increases in N fixation rates in our cold- and mesic-adapted BSC were caused by increases in cyanobacterial abundance, migration of cyanobacteria to the BSC surface, or a combination of both.

Between 20 and 25°C, we observed an increase in N fixation rates with no increase in cyanobacterial cover, suggesting that it may have been caused by changes in the metabolic activity of the N fixers. In a semi-arid *Pinus halepensis* plantation, warming did not affect the ratios between major microbial groups in lichen and moss crusts, but it did affect the level of physiological stress of the Gram negative bacterial community, as indicated by phospholipid fatty acid ratios (Maestre et al., 2015). Similarly, in soil from a temperate forest, short-term pulses of microbial respiration were caused by metabolic activation of dormant microbes, with no changes in total microbial biomass (Salazar-Villegas et al., 2016). Our observations suggest that under certain conditions the environment can affect N fixation rates in BSC by altering the metabolic rates of major N fixers, like cyanobacteria, even if there are no changes in their net abundance.

Much of the research on the effects of climate change on ecosystem function focuses on temperature and moisture (Zelikova et al., 1012; Hu et al., 2014; Salazar-Villegas et al., 2016; Rousk et al., 2018). The comparatively lower number of studies that have included light intensity have found, as in our own study, clear interactions between light, temperature and moisture on the cycling of elements through BSC (Lange et al., 1998; Grote et al., 2010). Interestingly, we did not find a relationship between N fixation rates and Chl *a*, as hypothesized. At the short temporal scale of our experiment, the effects of light on C fixation may be more dependent on the activity than on the abundance of photosynthetic cells and pigments. Because of the great dependence of N fixation on photosynthetically fixed C, the projected decreases in downward solar radiation in the high north by 2100 (as much as -10 W m^-2^ relative to the reference period of 1986-2005, under the RCP4.5 scenario; KNMI, 2021), could reduce N fixation rates in cold-adapted BSC.

The ways in which BSC interacts with the environment largely depends on its community composition (Belnap, 2002a; Bowker et al., 2021). On one hand, the rates at which BSC assimilates C and N depend on the abundance, type and activity of species in the system. BSC dominated by *Collema* soil lichens, for example, has higher levels of nitrogenase activity and therefore can fix more N than BSC dominated by *Microcoleus vaginatus* (Belnap, 2002a). On the other hand, the capacity of a BSC to resist environmental change depends on its community composition (Bowker et al., 2021). The presence of the lichens *Enchylium* and *Peltula* can increase the capacity of BSC to resist stress caused by wetting pulses at supra-optimal temperatures (Bowker et al., 2021).

It is well known that warming and moisture can induce significant compositional change in the bacterial communities of biocrust (Garcia-Pichel et al., 2013; Steven et al., 2015; Delgado-Baquerizo et al., 2018), with associated effects on bacterial functions (Steven et al., 2015). This can happen, for example, by environmental change differentially altering the growth rate and survival of different taxonomic groups (Lürling et al., 2013). Our results support the idea that warming can exert a selective pressure on warm-adapted microbial groups over those adapted to colder regimes (Muñoz-Martín et al., 2018). Moreover, our findings show that this selective pressure can manifest within a few days after environmental change. This was observed in our metagenomic investigations where photosynthesis-related genes are enriched at 10 °C but not at 20 °C, which corresponds to a decreased relative abundance of Nostocales. In addition, pathways related to a heterotrophic lifestyle are enriched at higher temperature and moisture, which correspond to a shift in the community composition with Alphaproteobacteria (e.g., Burkholderiales and Sphingomonadales) becoming more dominant in those conditions. In a broader sense, this suggests that climate change could lead to a replacement of the photosynthetic and N fixing Nostocales by non-photosynthetic N fixing bacteria. Such alteration could affect photosynthetic activity supported by cold adapted biocrust, and thus C cycling, as well as N cycling. A deeper understanding of the contribution of microbial community composition to Nfixation activity appears to be critical to predicting the productivity of cold biocrusts.

Short-term ecosystem responses to the environment may not necessarily reflect long-term trends (Alatalo et al., 2015). More multi-year, in situ experiments are needed at high latitudes (e.g. Rousk et al., 2018; Salazar et al., in progress) to better understand the effects of climate change on cold- and mesic-adapted BSC over large spatial and temporal scales. However, our short-term laboratory experiment 1) highlights direct and interacting effects of environmental factors on BSC composition and functioning, which could be useful to inform the structure and/or parameterizarion of ecosystem models that explicitly take into account the inherent dynamics of BSC (Rodríguez-Caballero et al., 2015); 2) provides data that can help with BSC characterization (Pietrasiak et al., 2013), which could further serve as reference when assessing the impacts of human disturbances on BSC (Szyja et al., 2018) and/or with the design and implementation of BSC-based restoration practices on degraded land (Bowker, 2007; Velasco-Ayuso et al., 2017; Tucker et al., 2020); and 3) contributes to the understanding of pulses of BSC activity in response to short-term climatic anomalies such as increasingly frequent and intense heat waves.

## Acknowledgments

Most AS work, including data collection and analysis, was done when working as a postdoc at University of Iceland, under the Icelandic Research Fund 2016 grant number 163336. AS finished his contributions to this article as an Assistant Professor at the Faculty of Environmental and Forests Sciences, Agricultural University of Iceland.

