## Supplementary information for "Environmental change alters nitrogen fixation rates and microbial parameters in a subarctic biological soil crust"

**Supporting information**

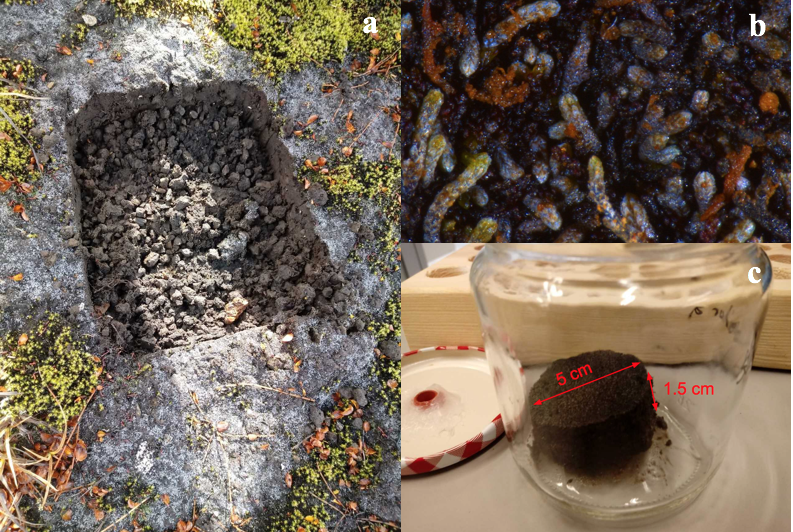

**Figure S1.** Photos of a) A landmark that illustrates the size of the eight BSC blocks (13x16 cm^2^, 5 cm depth) collected in the field. The photo shows the rocky vitrosol base below the BSC, and the moss (green) and vascular plants (brown) close to the BSC. The white color of the BSC surface comes from the wax on the leaves of the liverwort *A. juratzkana*; b) Stereoscope image (10x magnification) of a BSC sample; c) 5 cm diameter BSC disk used for measuring N fixation rates (section 2.3).

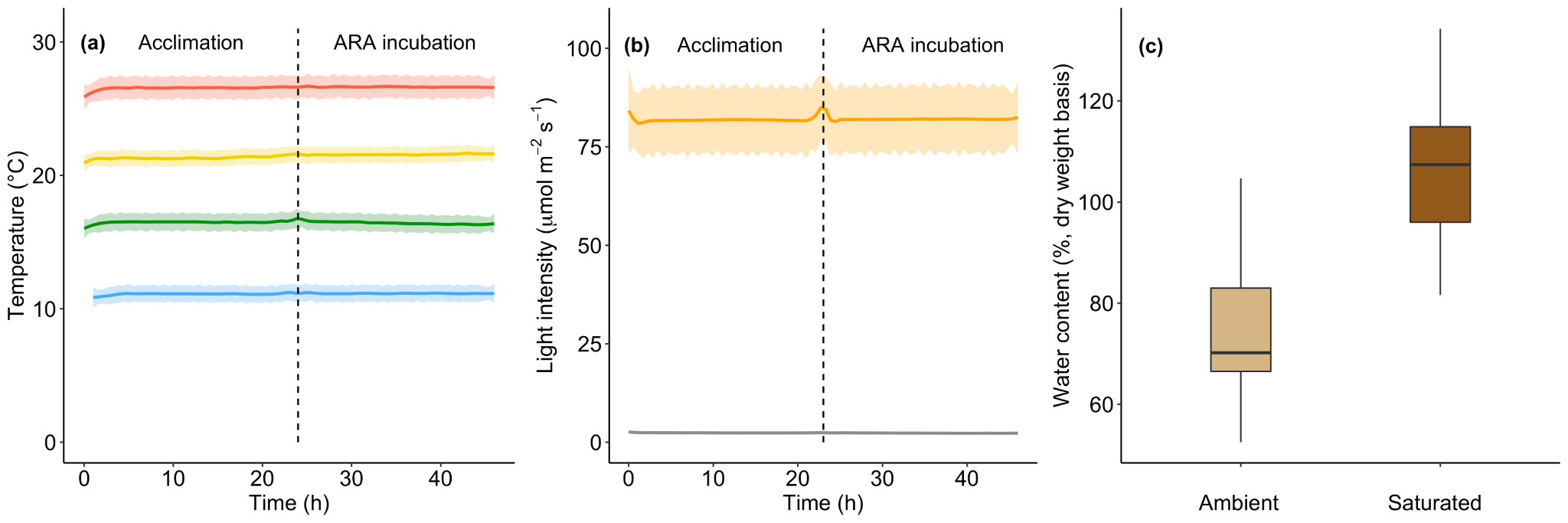

**Figure S2.** Levels of a) temperature, b) light intensity and c) moisture in our experimental design. a) Temperature levels 11.1 ± 0.10, 16.5 ± 0.10, 21.4 ± 0.10, and 26.6 ± 0.1 °C are shown in blue, green, yellow and red, respectively. b) Light intensity levels 2.3 ± 0.10 and 88.3 ± 1.0 μmol m^-2^ s^-1^ are shown in grey and orange respectively. c) Ambient (75.5 ± 2.4%) and at saturation (107.2 ± 2.3%) moisture content is shown in light and dark brown, respectively. The dashed line in a) and b) indicates the time when the acclimation period ended and the incubation period started (i.e. acetylene was injected). Values are means ± SE. n = 8.

-

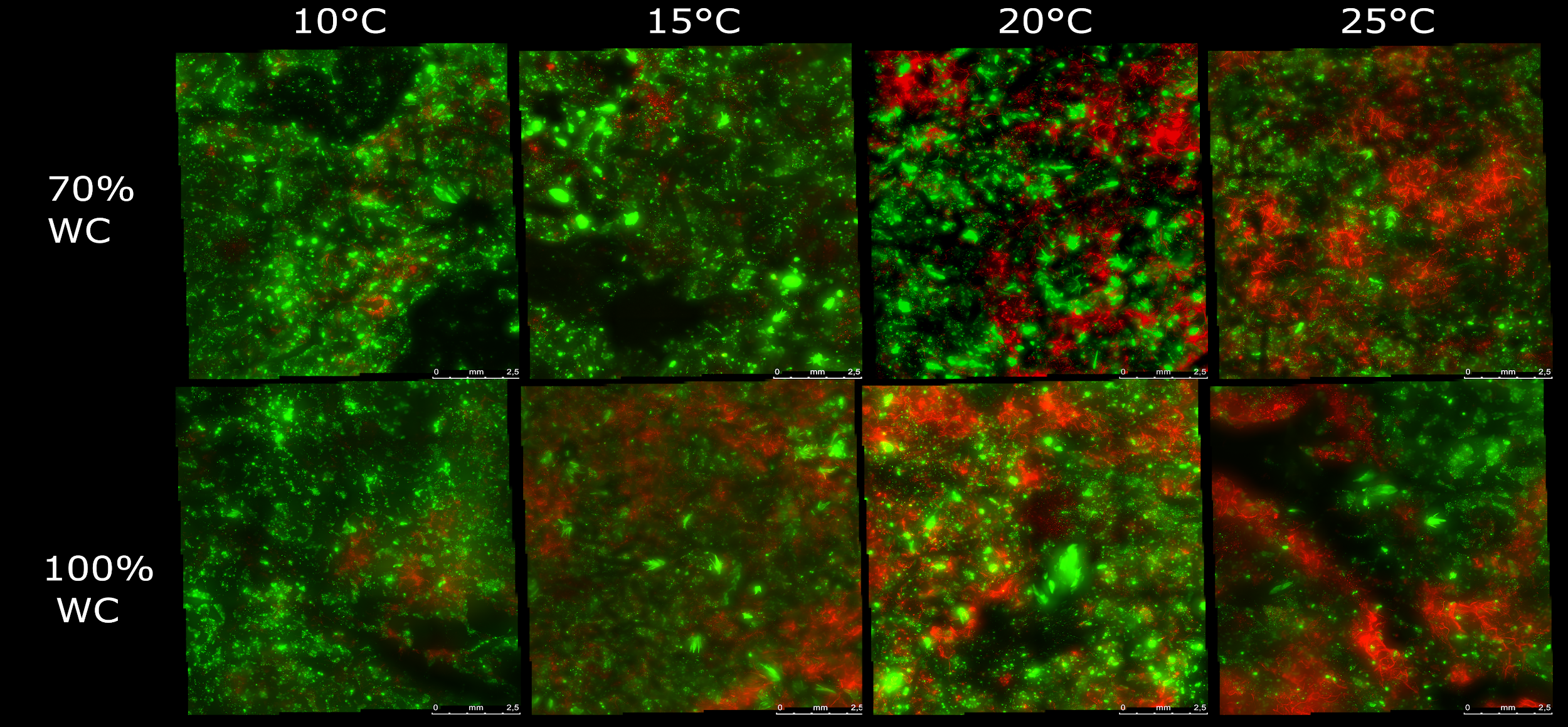

**Figure S3.** Epifluorescence microscopy images showing the cover of cyanobacteria (red) and liverwort (*Anthelia juratzkana*; green) on the surface of BSC after incubation at different levels of moisture and temperature for four days. Images were generated and analyzed using ImageJ/Fiji as described in the metho-ds.

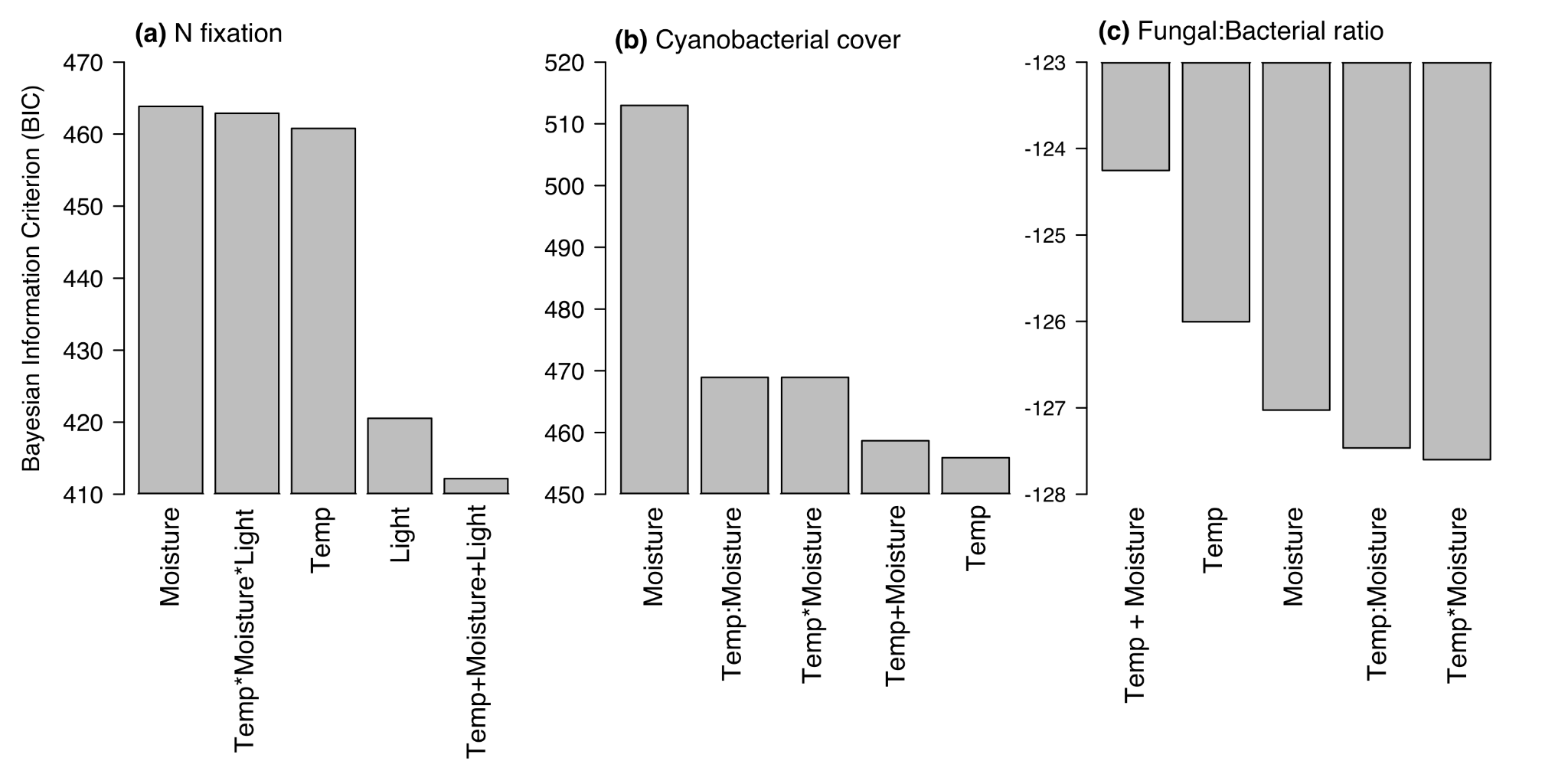

**Figure S4.** Comparison of statistical models of correlation based on the Bayesian Information Criterion (BIC) and R^2^ for a) N fixation rates, b) Cyanobacterial cover on BSC, and c) fungal:bacterial ratios. R^2^ is shown only when p<0.05. We compared models when at least one model was significant (i.e. no model comparisons for Chl *a*, total DNA or the relative abundance of cyanobacterial groups).

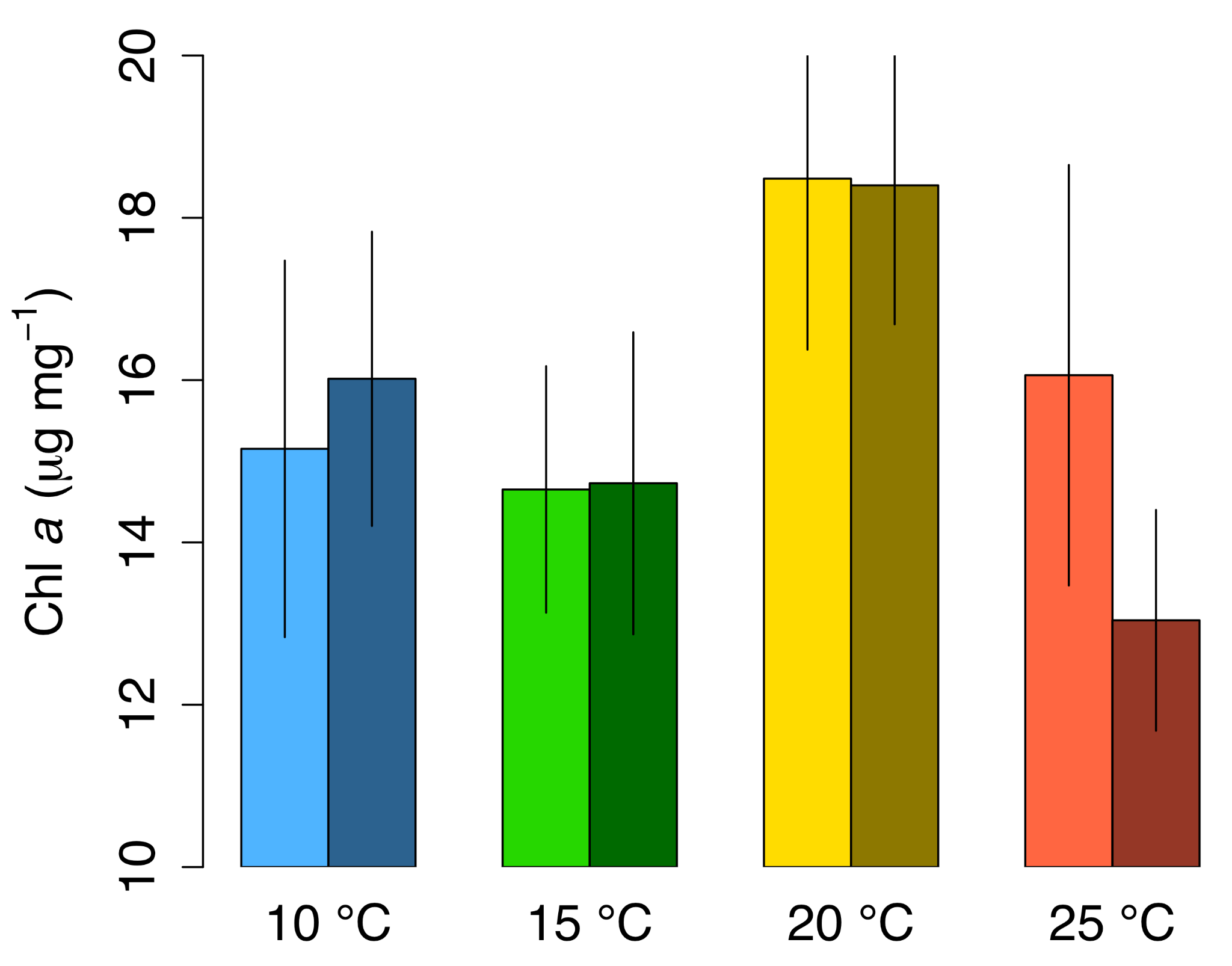

**Figure S5.** Chl *a* content of BSC under manipulated temperatures and moistures. Blue, green, yellow and red colors indicate 10, 15 and 20 and 25 °C, and light and dark colors indicate 75% and saturated moisture content, respectively. Values are mean ± SE. Averages are from eight samples, combined into four (n=4). There were no differences (P > 0.05) in Chl *a* content between treatments.

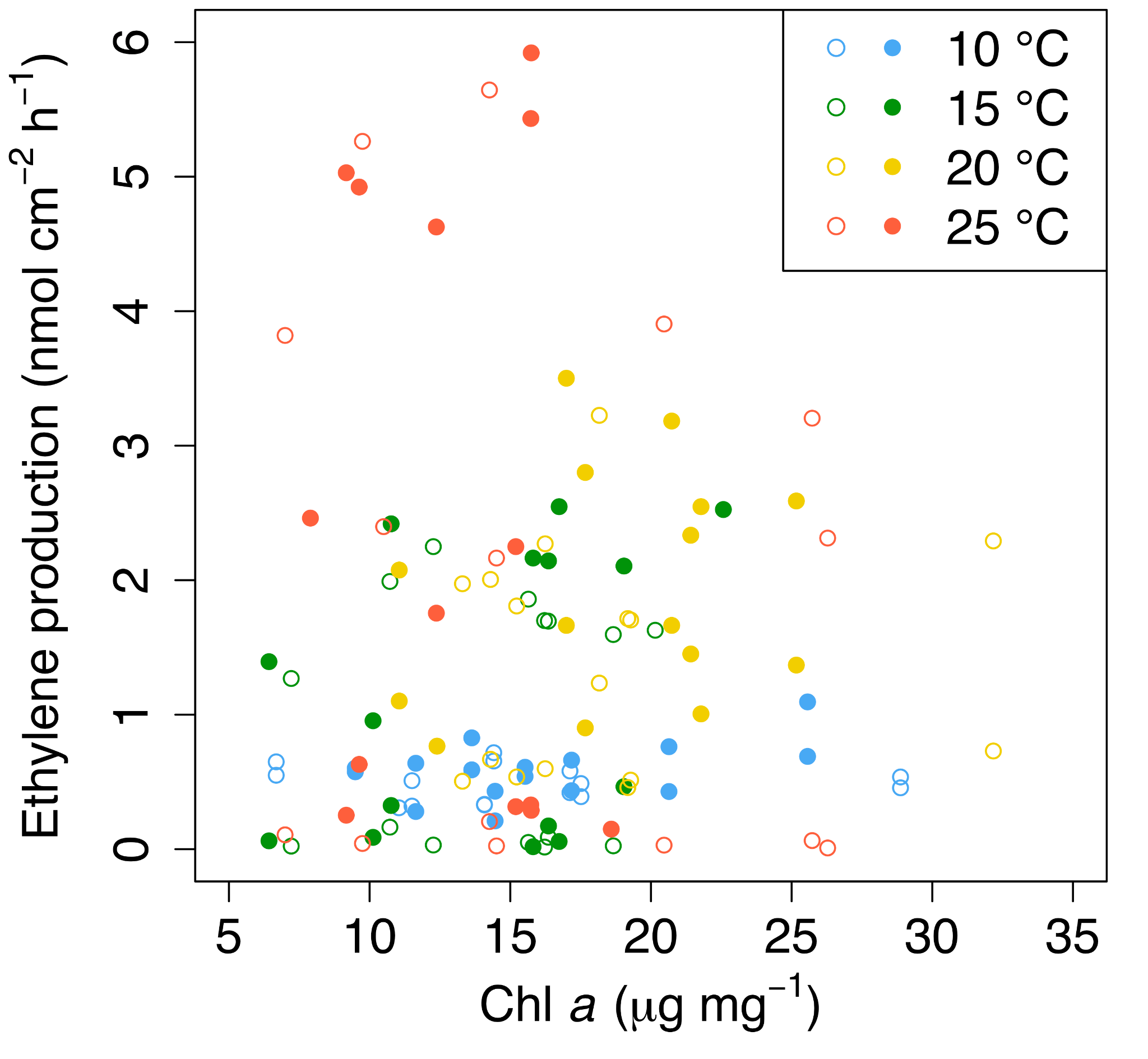

**Figure S6.** Lack of correlation (P > 0.05) between ethylene production and Chl *a* content in BSC under different temperature and moisture treatments. There are two values of ethylene production per value of Chl *a*, one at ca. 2 μmol m^-2^ s^-1^ of light, and the other at ca. 90 μmol m^-2^ s^-1^.

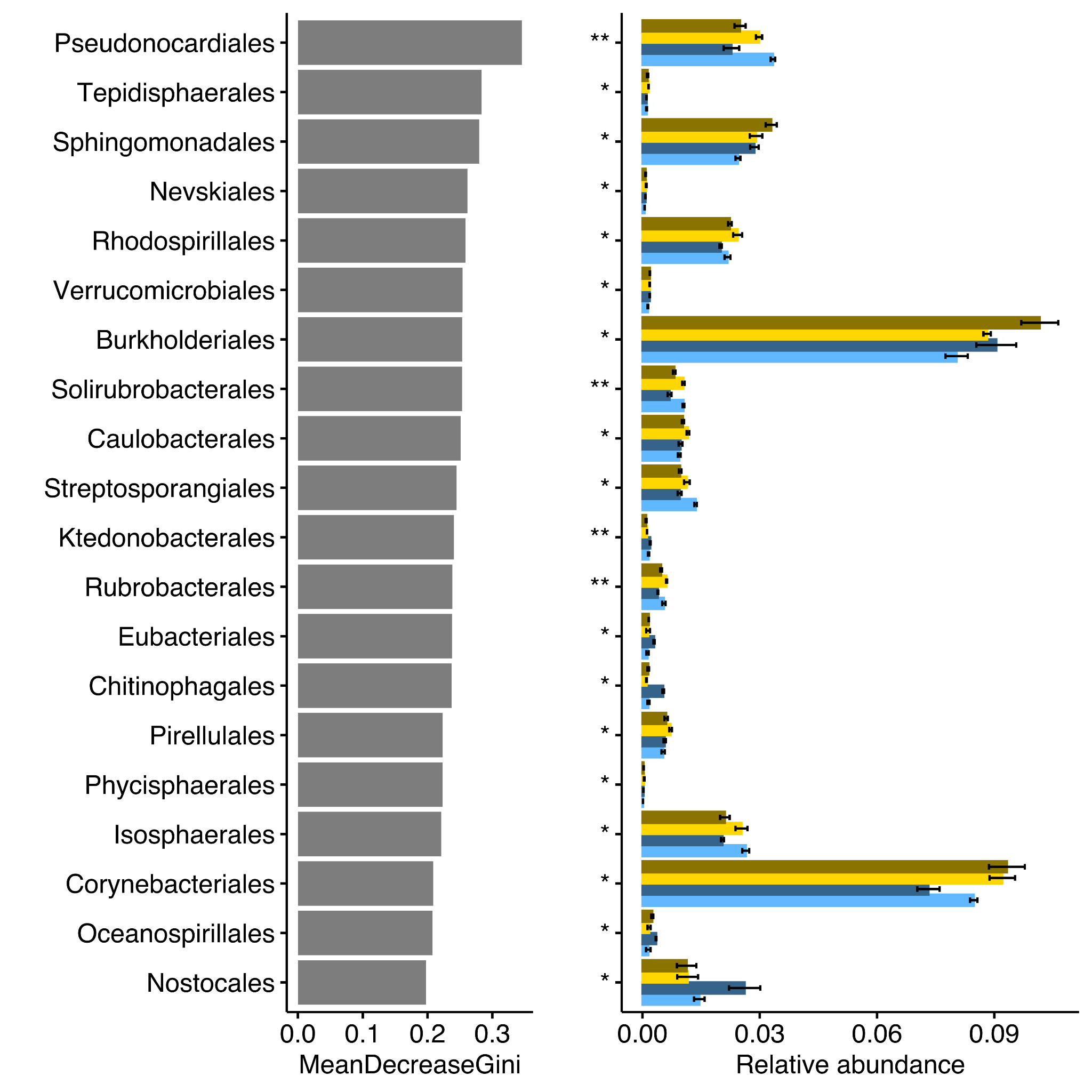

**Figure S7.** Top 20 important Orders associated with changes in moisture and temperature identified by Random Forest classification. Top 20 taxa were assessed by Gini index, which represented the importance of each Order in distinguishing different groups. The read abundances of the identified top 20 taxa are shown on the right panel. Blue and yellow color indicate 10 and 20 °C, and light and dark tones indicate 70 and 100% moisture content, respectively. Significant Gini index after 100 bootstraps at *, p< 0.05; **, p< 0.01; ***, p< 0.001.

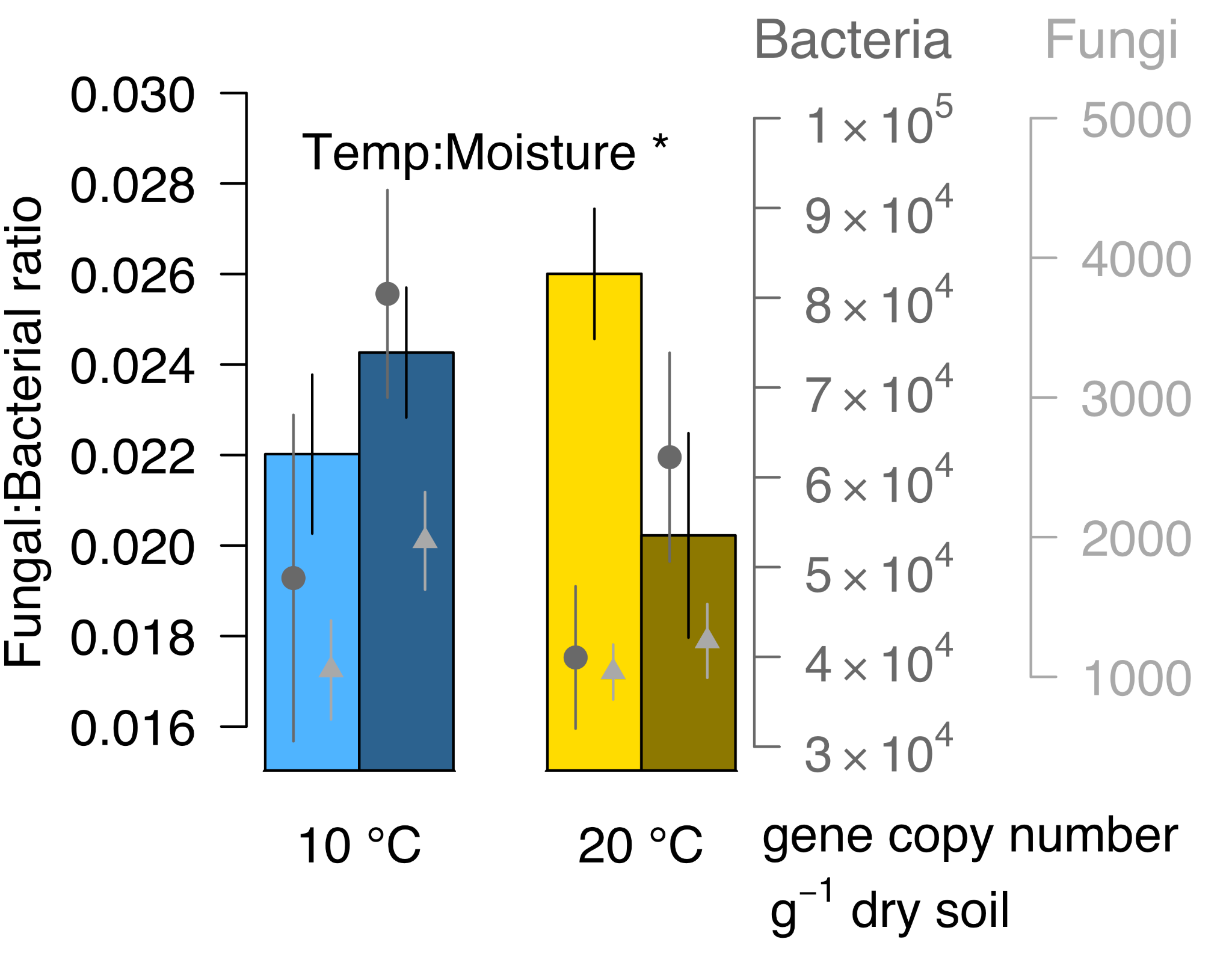

**Figure S8.** Fungal:bacterial gene ratios (bars and black y axis on the left) based on gene copy number from bacteria (dark grey circle marks and axis on the right) and fungi (light grey triangle marks and axis on the right) as determined by Kaiju analysis. Blue and yellow colors indicate 10 and 20 °C, and light and dark bars indicate 70 and 100% moisture content, respectively. Gene copy numbers for bacteria and fungi are normalized by the amount of DNA extracted per gram of dry sample. Values are mean ± SE. Averages are from eight samples, combined into four (n=4).

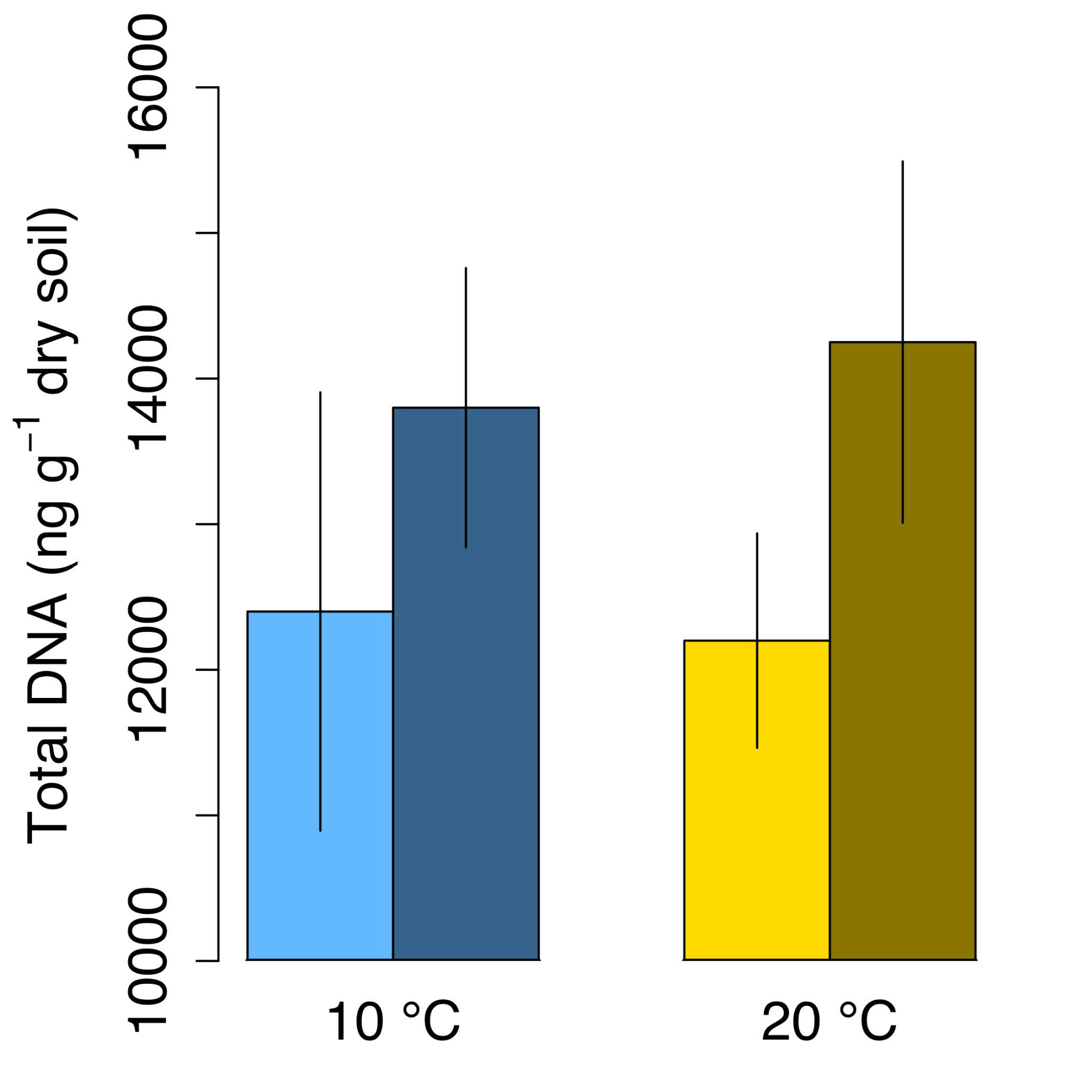

**Figure S8.** Total DNA measured with a Qubit 4 Fluorometer (Thermo Fisher Scientific, USA). Blue and yellow colors indicate 10 and 20 °C, and light and dark colors indicate 70 and 100% moisture content, respectively. Values are mean ± SE. Averages are from eight samples, combined into four (n=4). There were no significant differences in total DNA between treatments.

**Table S1.** ADONIS results based on Bray-Curtis distance for the response of microbial communities composition to the simulated temperature and moisture treatments on soil biocrust.

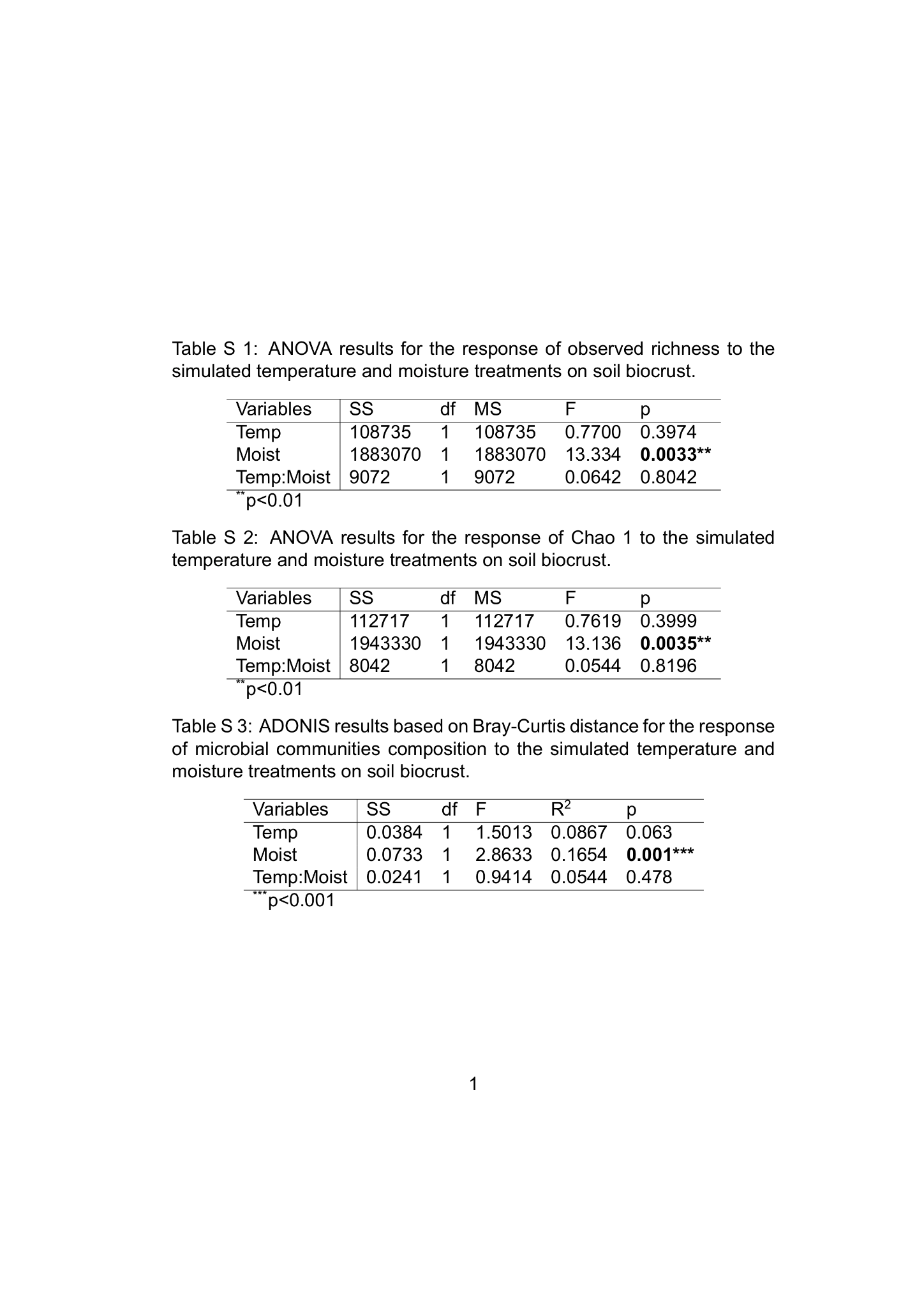

**Table S2.** ANOVA results for the response of observed richness to the simulated temperature and moisture treatments on soil biocrust.

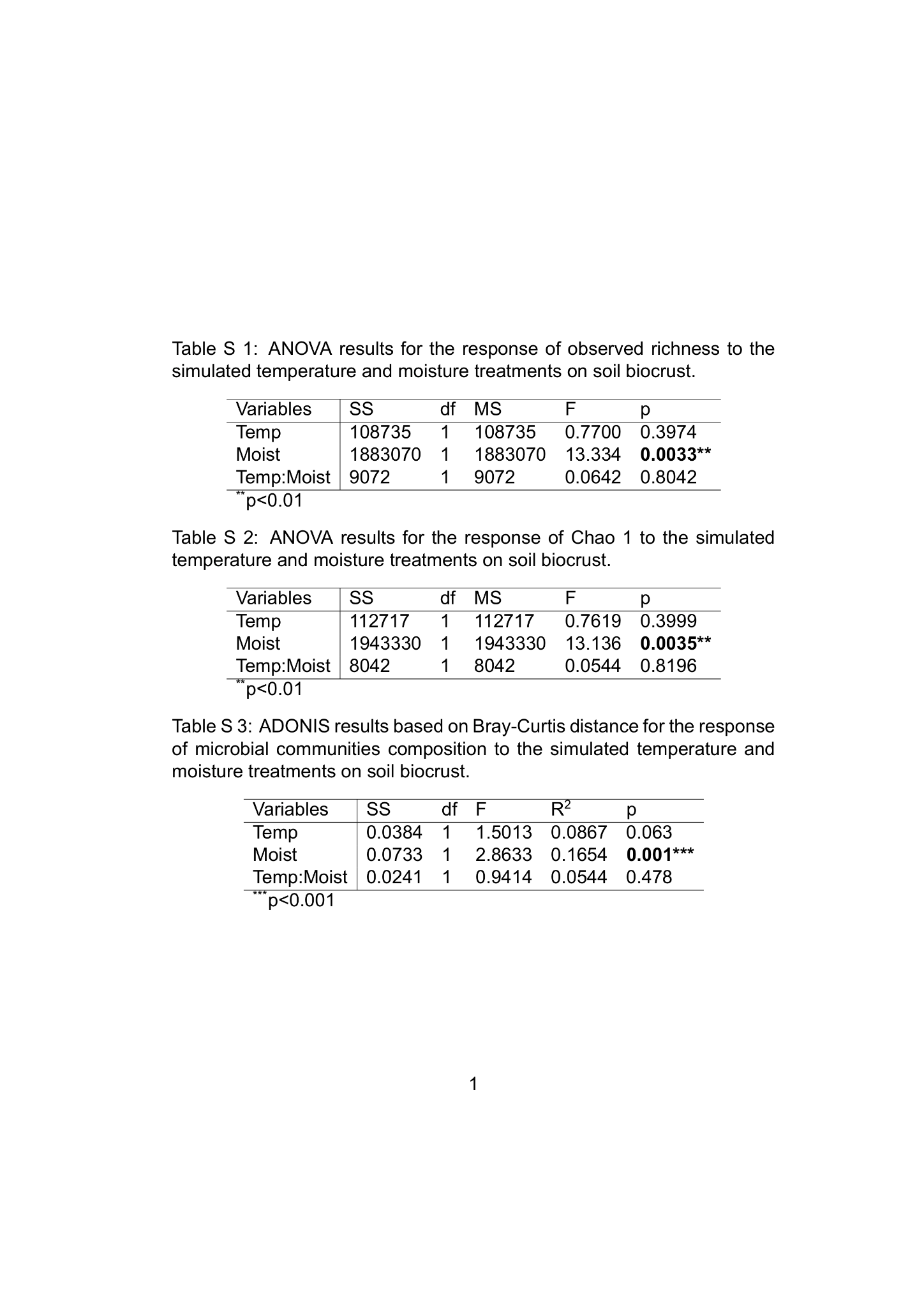

**Table S3.** ANOVA results for the response of Chao 1 to the simulated temperature and moisture treatments on soil biocrust.

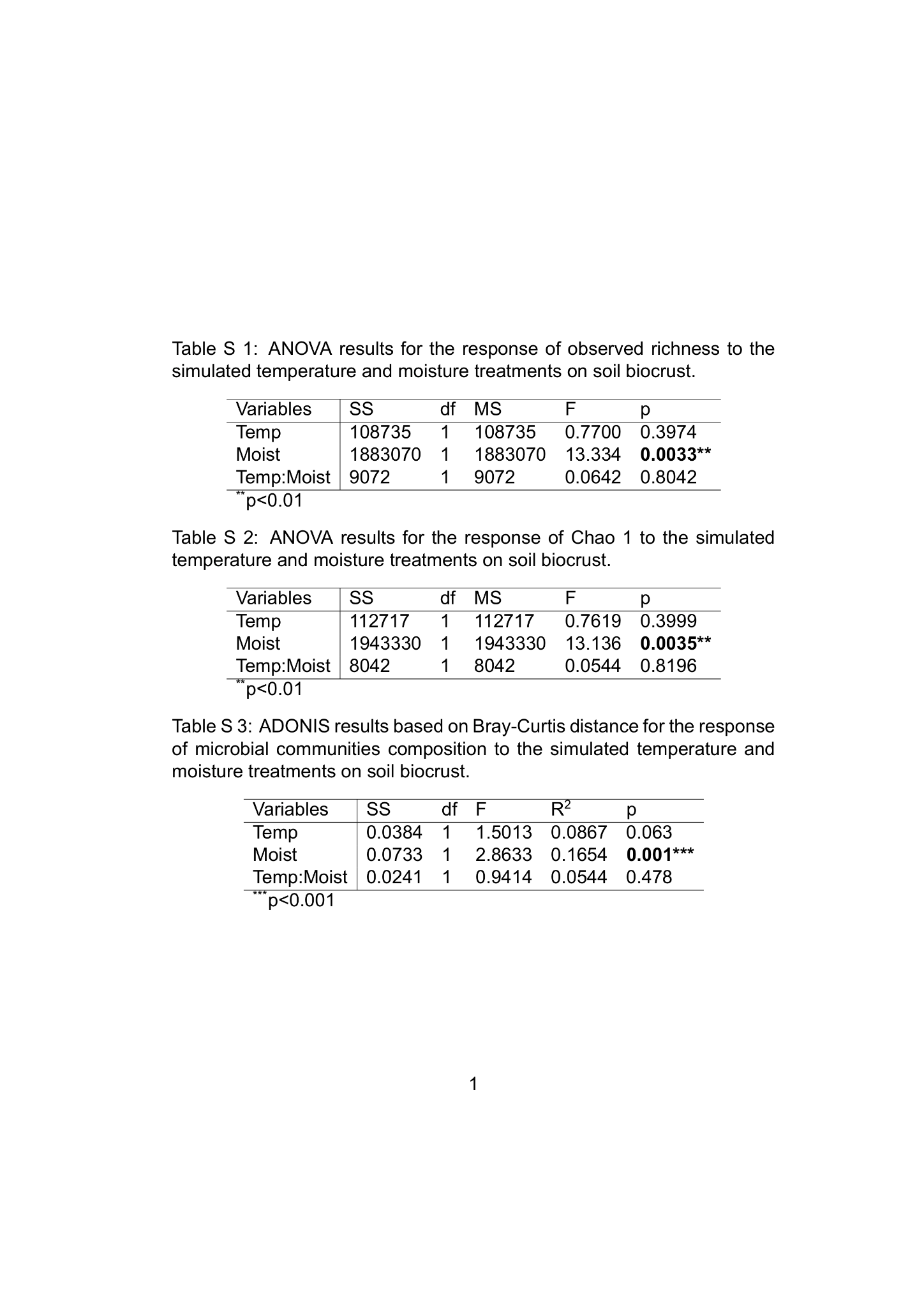
